# Directional coupling of slow and fast hippocampal gamma with neocortical alpha/beta oscillations in human episodic memory

**DOI:** 10.1101/305698

**Authors:** Benjamin J. Griffiths, George Parish, Frederic Roux, Sebastian Michelmann, Mircea van der Plas, Luca D. Kolibius, Ramesh Chelvarajah, David T. Rollings, Vijay Sawlani, Hajo Hamer, Stephanie Gollwitzer, Gernot Kreiselmeyer, Bernhard Staresina, Maria Wimber, Simon Hanslmayr

## Abstract

Episodic memories hinge upon our ability to process a wide range of multisensory information and bind this information into a coherent, memorable representation. On a neural level, these two processes are thought to be supported by neocortical alpha/beta desynchronisation and hippocampal theta/gamma synchronisation, respectively. Intuitively, these two processes should couple to successfully create and retrieve episodic memories, yet this hypothesis has not been tested empirically. We address this by analysing human intracranial EEG data recorded during two associative memory tasks. We find that neocortical alpha/beta (8-20Hz) power decreases reliably precede and predict hippocampal “fast” gamma (60-80Hz) power increases during episodic memory formation; during episodic memory retrieval however, hippocampal “slow” gamma (40-50Hz) power increases reliably precede and predict later neocortical alpha/beta power decreases. We speculate that this coupling reflects the flow of information from neocortex to hippocampus during memory formation, and hippocampal pattern completion inducing information reinstatement in the neocortex during memory retrieval.

**Significance Statement:** Episodic memories detail our personally-experienced past. The formation and retrieval of these memories has long been thought to be supported by a division of labour between the neocortex and the hippocampus, where the former processes event-related information and the latter binds this information together. However, it remains unclear how the two regions interact. We uncover directional coupling between these regions, with power decreases in the neocortex that precede and predict power increases in the hippocampus during memory formation. Fascinatingly, this process reverses during memory retrieval, with hippocampal power increases preceding and predicting neocortical power decreases. These results suggest a bidirectional flow of information between the neocortex and hippocampus is fundamental to the formation and retrieval of episodic memories.

## Introduction

An episodic memory is a high-detailed memory of a personally-experienced event^1,2^. The formation and retrieval of such memories hinge upon: a) the processing of information relevant to the event, and b) the binding of this information into a coherent episode. A recent framework^3^ and computational model^4^ suggest that the former of these processes is facilitated by the desynchronisation of neocortical alpha/beta oscillatory networks (8-20Hz; reflected in decreases in oscillatory power)^5^, while the latter is facilitated by the synchronisation of hippocampal theta and gamma oscillations (3-7Hz; 40-100Hz; reflected in increases in oscillatory power)^6,7^ [see fig. 1a]. Critically, the framework posits that these two mechanisms need to cooperate, as an isolated failure of either of these mechanisms would produce the same undesirable outcome: an incomplete memory trace. Here, we test this framework and provide the first empirical evidence of an interaction between neocortical desynchronisation and hippocampal synchronisation during the formation and retrieval of human episodic memories. In addition, we demonstrate that distinct hippocampal gamma frequencies contribute to memory formation and retrieval, with “fast” gamma facilitating encoding and “slow” gamma facilitating retrieval.

**Figure 1.**
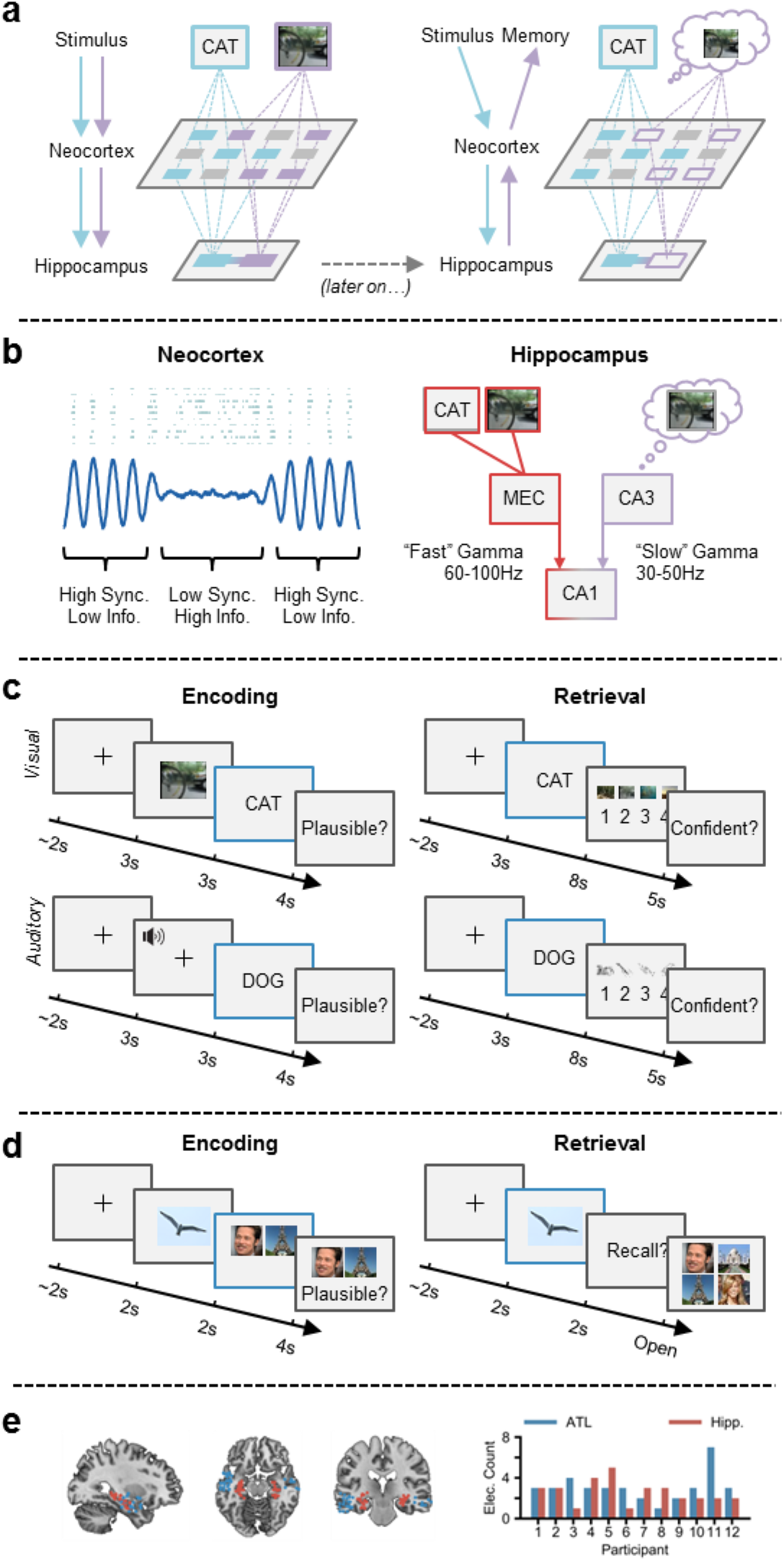
The sync-desync framework. **(a)** Incoming stimuli are independently processed by relevant sensory regions of the neocortex (left), and then passed onto the hippocampus where they are bound together. At a later stage (right), a partial cue reactivates the hippocampal associative link, which in turn reactivates neocortical patterns coding for the memory representation, giving rise to conscious recollection. **(b)** Reduced oscillatory synchronisation (blue line) within the neocortex allows individual neurons (blue dots) to fire more freely and create a more flexible neural code. “Fast” gamma activity allows from the transferal of neocortical information to the hippocampus by boosting connectivity between the entorhinal cortex (MEC) and CA1. “Slow” gamma enhances retrieval by boosting connectivity between CA3 and CA1, allowing reinstated memories to be passed to the neocortex. **(c)** During encoding, participants are tasked with forming an associative link between a life-like dynamic stimulus (either a video or sound) and a subsequent verbal stimulus. During retrieval, participants are presented with verbal stimuli from the previous encoding block and asked to retrieve the associated dynamic stimulus. Electrophysiological analysis was conducted during the presentation of the verbal stimulus at encoding and retrieval (blue outline). **(d)** During encoding, participants are tasked with forming an associative link between an object, a face and a scene. During retrieval, participants are presented with the object and asked to retrieve the associated face and scene. Electrophysiological analysis was conducted during the presentation of the verbal stimulus at encoding and retrieval (blue outline). **(e)** Plot of each electrode location (left; red represents hippocampal electrode; blue represents ATL. Bar plot (right) depicts number of electrodes for each participant.

Within the neocortex, desynchronised alpha/beta activity is thought to facilitate information processing^5^. This hypothesis is based on the principles of information theory^8^, which proposes that a system of unpredictable states (e.g. desynchronised neural activity, where the firing of one neuron is not predictive of the firing of another; see Hanslmayr et al., 2012 for details) is optimal for information coding (see fig. 1b). Neural desynchronisation in humans is most often measured by a decrease in oscillatory power, as a strong correlation exists between neural synchronisation and power^9^ (though this link is strictly correlative). In support of the information-via-desynchronisation hypothesis, many studies have observed neocortical alpha/beta power decreases during successful episodic memory formation^10–18^ and retrieval^19–24^. For example, neocortical alpha/beta power decreases scale with the depth of semantic processing during episodic memory formation^18^. Critically, synchronising alpha/beta rhythms via repetitive transcranial magnetic stimulation impairs both episodic memory formation and retrieval, suggesting that alpha/beta desynchronisation plays a causal role in these processes^20,25^. In conjunction, these studies suggest that neocortical alpha/beta desynchronisation underpins the processing of event-related information, all owing for the formation and later recollection of highly detailed episodic memories.

Within the hippocampus, synchronised gamma activity (30-100Hz) is thought to be critical in the binding of event-related information, and the later retrieval of this information when prompted by a cue^6,7,26,27^. Entraining neurons to rhythms of approximately 60Hz (i.e. a “fast” gamma oscillation) allows for spike-timing dependent plasticity (STDP; a form of long-term potentiation) to occur^28^, which strengthens synaptic connections between hippocampal neurons. As such, an increase in hippocampal “fast” gamma activity (60-100Hz) may be a proxy for STDP^29,30^ and, therefore, representational binding. In contrast, a slower hippocampal gamma rhythm (30-50Hz) has been proposed to facilitate memory retrieval^7,31,32^. “Slow” gamma activity originates from the CA3 subfield of the hippocampus and may play a pivotal role in pattern completion^33,34^. The trade-off in amplitude between these two gamma oscillations is thought to dictate whether encoding or retrieval takes place^35^. Evidence suggests that periods of increased “fast” gamma activity enhances connectivity between CA1 and the entorhinal cortex^31,36^ (allowing information to flow into the hippocampus; see fig. 1b) and aids representational binding through STDP^28,30^. Meanwhile, periods of enhanced “slow” gamma activity sees an increase in connectivity between CA1 and CA3 (allowing for the transfer of completed memory pattern into the neocortex; see fig. 1b)^31,36^. In conjunction, these findings and theories would suggest that “fast” and “slow” gamma rhythms differentially support the hippocampal ability to associate and reactivate discrete elements of an episodic memory.

Here, we investigated the co-ordination between alpha/beta power decreases in the anterior temporal lobe (ATL) and gamma power increases in the hippocampus during episodic memory formation and retrieval. Specifically, we tested four central hypotheses derived from a series of conceptual frameworks, computational models and rodent studies: 1) “fast” gamma oscillations (60-100Hz) will support encoding while “slow” gamma oscillations (30-45Hz) will support memory retrieval^7,31^; 2) neocortical power decreases (reflecting information processing^5^) and hippocampal power increases (reflecting representational binding^6,7,26,27^) will accompany episodic memory formation and retrieval when contrasted against memories that were not successfully encoded/retrieved; 3) neocortical power decreases will precede hippocampal power increases during memory formation (reflecting information processing preceding representational binding), and hippocampal power increases will precede neocortical power decreases during retrieval (reflecting pattern completion preceding information reinstatement)^3,4^.

Twelve patients implanted with stereotactic EEG electrodes for the treatment of medication-resistant epilepsy completed one of two associative memory tasks (see fig. 1c-d; n=7 in task 1; n=5 in task 2). In task 1, they related life-like videos or sounds to words that followed. Following a short distractor task, participants attempted to recall the previously presented videos/sounds using the words as cues. In task 2, they related an object to pairs of visual stimuli that followed (face-place, face-face or place-place). Following a short distractor task, participants attempted to recall both stimuli, using the object as a cue. While external stimulation is different between the two tasks, the underlying cognitive and neural processes relating to our hypotheses are consistent: both tasks require sensory processing followed by representational binding during memory formation, and hippocampal pattern competition prior to neocortical reinstatement during memory retrieval. As such, the data from the two tasks were pooled together for analysis. We conducted these analyses in two ROIs (see fig. 1e): the hippocampus (a hub for representation binding) and the anterior temporal lobe (ATL; a hub for semantic-based information processing^37^). Foreshadowing the results below, we show that ATL alpha/beta power decreases precede hippocampal “fast” gamma power increases during successful memory formation, and that hippocampal “slow” gamma power increases precede ATL alpha/beta power decreases during successful memory retrieval. The results reveal the first empirical evidence of an interaction between these two oscillatory dynamics during human episodic memory formation and retrieval.

## Results

### Behavioural results

Participants, on average, recalled 47.9% of all pairs in the first task, a percentage much greater than what would be expected by chance (25%). When breaking trials down by modality, participants recalled 52.7% of video-word pairs and 45.9% of sound-word pairs. An independent samples t-test (only a subset of participants completed both variants of the task) revealed no significant difference in memory performance for video-word and sound-word pairs (p > 0.5, d = 0.275). As there was no apparent difference in memory performance between the two trials types, and electrode contacts were not located in anatomical regions that should respond uniquely to one of these sensory modalities, trials involving video-word and sound-word pairs were combined for all further analyses. In the second task, participants recalled both associated items on 66.2% of trials - a percentage much greater than what would be expected by chance (16.7%; where the probability of selecting the first item correctly is 50% and the probability of selecting the second item correctly is 33%, making the joint probability 50% × 33% = 16.7%).

### Distinct oscillatory signatures exist in the neocortex and hippocampus

We first sought to empirically define the peak frequencies in our three regions of interest. Broadband spectral power (1-100Hz) was computed using 5-cycle wavelets across a 1500ms window starting at the onset of the verbal stimulus (at encoding and retrieval). The data was then z-transformed across trials to facilitate comparison across participants, and the 1/f component was subtracted from the data^38–40^ to attenuate broadband noise (see methods for details). Subsequently, the resulting power spectra were collapsed over time and trials, and split into hippocampal and neocortical ROIs. Across participants, a slow-theta peak could be observed in the hippocampus at ∼2.5Hz and an alpha/beta peak could be observed in the two neocortical regions between 8-20Hz (see figure 2a). We defined the peak frequency of each ROI for each participant individually and conducted all subsequent analyses on these peak frequencies (see SI appendix, table S1 for individual peak frequencies).

**Figure 2.**
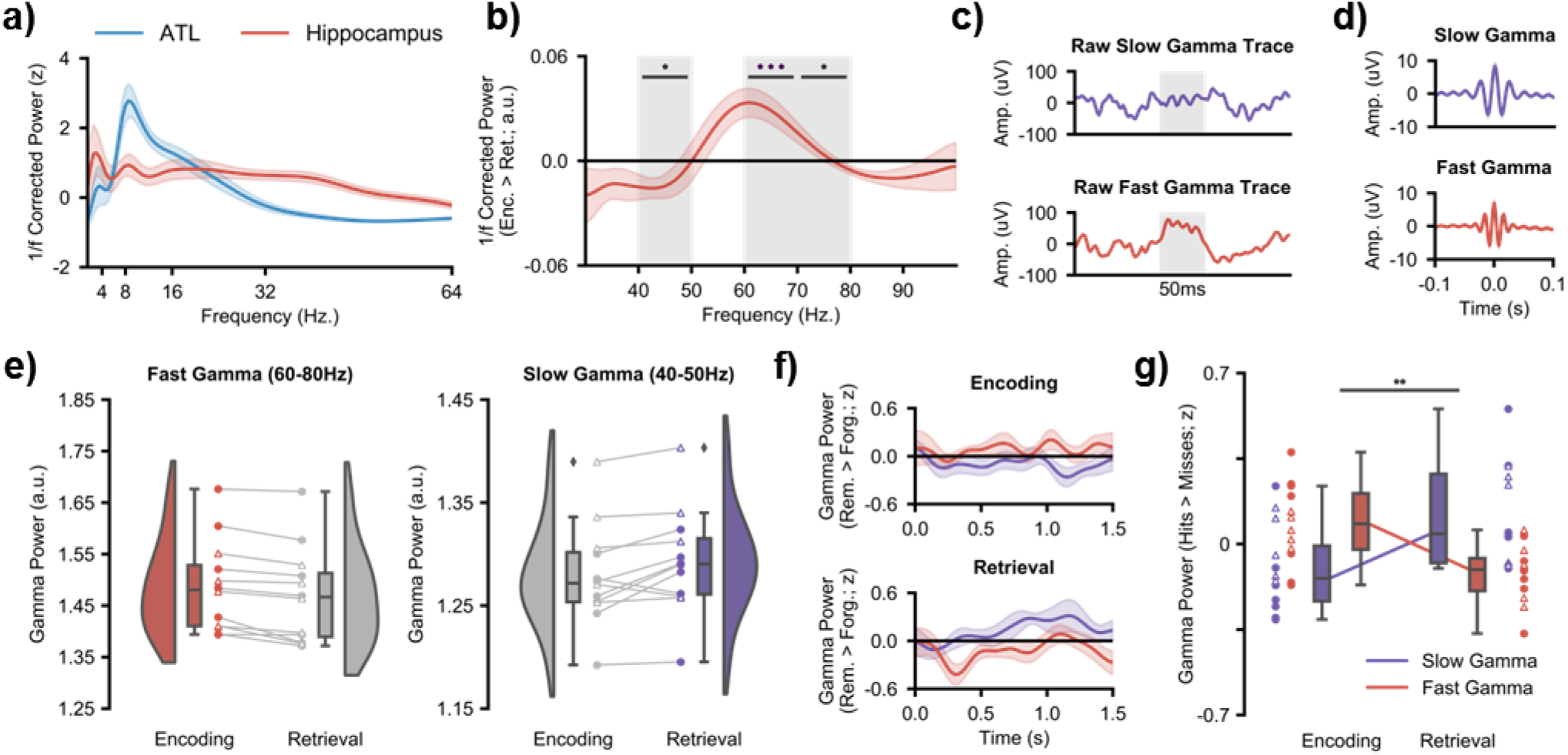
Hippocampal gamma activity during encoding and retrieval. **(a)** the mean 1/f corrected power spectrum (with shaded standard error of the mean) across all encoding and retrieval trials reveals theta and gamma peaks in the hippocampus and an alpha/beta peak peak in the ATL. **(b)** the mean difference in gamma power (with shaded standard error of the mean) between encoding and retrieval reveals a peak in encoding-related, “fast” gamma at 60-80Hz and a peak in retrieval-related, “slow” gamma at 40-50Hz (*p_fdr_ < 0.05, ***p_fdr_ < 0.001. **(c)** raw slow gamma signal during retrieval (top) and fast gamma signal during encoding (bottom) from a hippocampal contact of participant 1. The shaded grey region indicates a period of 50 milliseconds. **(d)** mean peak-locked averaged signal across participants for slow (top) and fast (bottom) gamma (with shaded standard error of the mean). **(e)** raincloud plots depicting the difference in fast (left) and slow (right) gamma power between encoding and retrieval. Coloured circles represent participants who took part in experiment 1. Uncoloured triangles represent participants who took part in experiment 2. **(f)** time-series of slow (in purple) and fast (in red) memory-related gamma power for encoding and retrieval. **(g)** interaction between fast and slow gamma activity during encoding and retrieval. Encoding sees a relative increase of memory-related fast gamma power, while retrieval sees a relative increase of memory-related slow gamma power.

### Distinct hippocampal gamma-band frequencies underlie encoding and retrieval processes

We then investigated whether distinct gamma frequency bands support encoding and retrieval processes^7,31^. To test this, the broadband hippocampal gamma power (30-100Hz) for successfully remembered pairs at encoding and retrieval was calculated and contrasted in a group level, non-parametric permutation test. “Fast” hippocampal gamma frequencies (60-80Hz) exhibited significantly greater power during encoding, relative to retrieval, trials (60-70Hz, p_fdr_ = 0.001, d = 1.308; 70-80Hz, p_fdr_ = 0.020, d = 0.947; see fig. 2b-e). In contrast, “slow” hippocampal gamma frequencies (40-50Hz) exhibited greater power during retrieval, relative to encoding, trials (p_fdr_ = 0.023, d = 0.754). No significant difference between encoding and retrieval could be observed during the epochs of forgotten stimuli (see SI appendix, Fig. S1). Peak “fast” and “slow” gamma frequencies for each participant were derived from the “encoding vs. retrieval” contrast and used in all subsequent analyses (see methods for details; see SI appendix, Table S1 for individual peak frequencies). These findings provide the first empirical evidence that two functionally-relevant gamma band oscillations relate to episodic memory formation and retrieval in humans.

To rule out the possibility that the difference in “fast”/”slow” gamma was driven by the 1/f slope and/or its removal, the beta weights describing the 1/f slope at encoding and retrieval were extracted and averaged across time, electrodes and trials. These weights were then contrasted between encoding and retrieval in a group level, non-parametric permutation test. This test revealed no significant difference in the beta weights for remembered items (p = 0.198) or for forgotten items (p = 0.246), suggesting the distinction in gamma rhythms between encoding and retrieval was not driven by differences in the 1/f slope.

### Hippocampal gamma power increases track the successful formation and retrieval of episodic memories

To examine how memory-related fluctuations in “fast” and “slow” gamma power differentially contribute to episodic memory encoding and retrieval, we conducted a group level, non-parametric, permutation-based, 2×2 repeated measures ANOVA that investigated the influence of factors ‘gamma frequency’ (“fast” vs. “slow”) and ‘memory operation’ (encoding vs. retrieval) on memory-related power (remembered > forgotten) collapsed across time. We anticipated an interaction whereby “fast” gamma selectively supports successful memory formation and “slow” gamma selectively supports successful memory retrieval. Group analysis revealed a significant interaction (p = 0.003, partial η^2^ = 0.294; see figure 4g), indicating that “fast” and “slow” gamma exhibited dissimilar memory-related power fluctuations during encoding and retrieval. These results demonstrate that two functionally-distinct gamma band oscillations support the successful formation and retrieval of episodic memories in humans.

Analysis of the power time series showed that the opposing effects of “fast” and “slow” gamma was particularly prominent during retrieval. When successfully recalling a stimulus, a rapid decrease in “fast” gamma power was observed (200-400ms, p_fdr_ = 0.025, d = 0.862, see fig. 4f), followed by an increase in “slow” gamma power (800-1000ms, p_fdr_ = 0.007, d = 1.177, see fig. 4f), relative to stimuli that were not recalled. Perplexingly, a similar effect was not observed during encoding even though the time series of the two gamma bands trend in the correct directions (i.e. an increase in “fast” gamma and a decrease in “slow” gamma; see fig. 4f). As will be revealed later, this absence may be driven by the fact that gamma power changes are not time-locked to stimulus onset during encoding, but rather the neocortical power decreases that precedes hippocampal activity.

### Neocortical alpha/beta power decreases track the successful formation and retrieval of episodic memories

We then investigated whether neocortical alpha/beta power decreases accompany the successful encoding and retrieval of episodic memories. Peak alpha/beta power was computed across a 1500ms window commencing at stimulus onset. As above, the 1/f characteristic was subtracted, attenuating broadband noise^42^. The alpha/beta power was z-transformed across the entire session for each electrode-frequency pair separately, smoothed to attenuate trial-by-trial variability in temporal/spectral responses (see methods), and split into “hits” and “misses” for contrasting. A group level, non-parametric permutation test revealed a significant decrease in anterior temporal lobe (ATL) alpha/beta power during encoding (p_fdr_ = 0.035, d = 0.858; 400-600ms after stimulus onset, fig. 3) for remembered stimuli relative to forgotten stimuli. During retrieval, a group level, permutation test revealed a significant decrease in ATL alpha/beta power (800-1000ms, p_fdr_ = 0.042, d = 0.777; 1000-1200ms, p_fdr_ = 0.039, d = 0.849; see fig. 3) for remembered stimuli relative to forgotten stimuli. These results reproduce earlier findings of neocortical alpha/beta power decreases during the encoding^10–18^ and retrieval^19–24^ of human episodic memories.

**Figure 3.**
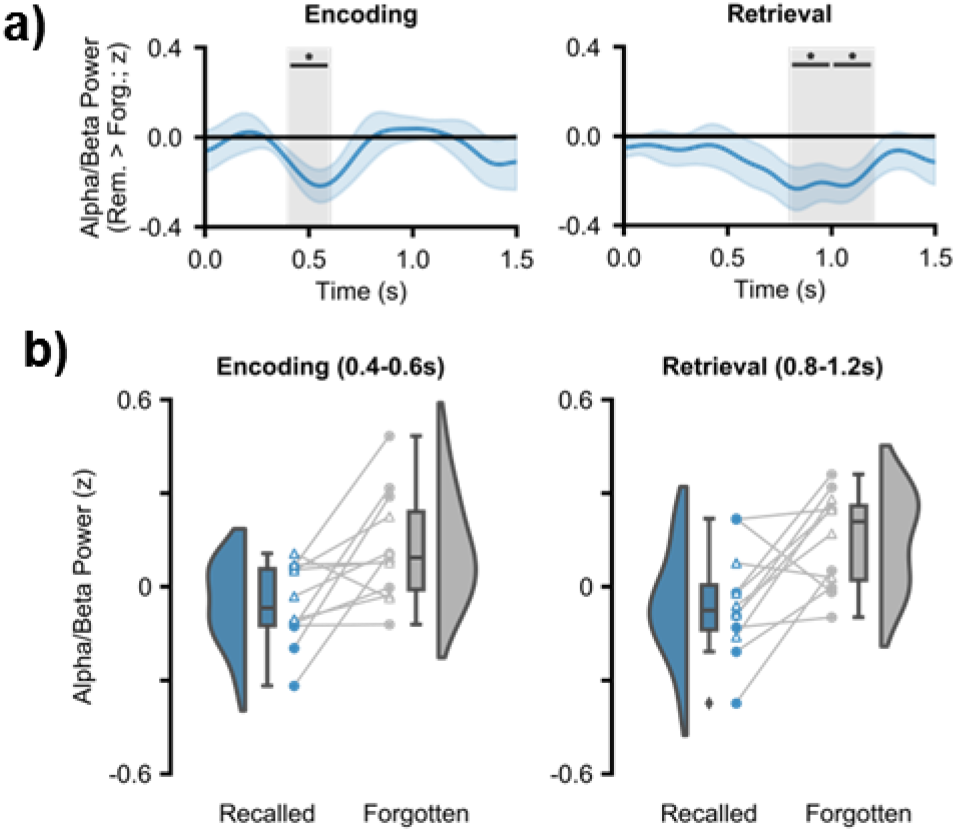
ATL alpha/beta activity during encoding and retrieval, **(a)** time-series of memory-related alpha/beta power for encoding and retrieval. In both cases, decreases in alpha/beta power relate to greater memory (*P_fdr_ < 0.05). **(b)** raincloud plots depicting the difference in alpha/beta power between remembered and forgotten items. Coloured circles represent participants who took part in experiment 1. Uncoloured triangles represent participants who took part in experiment 2.

**Figure 4.**
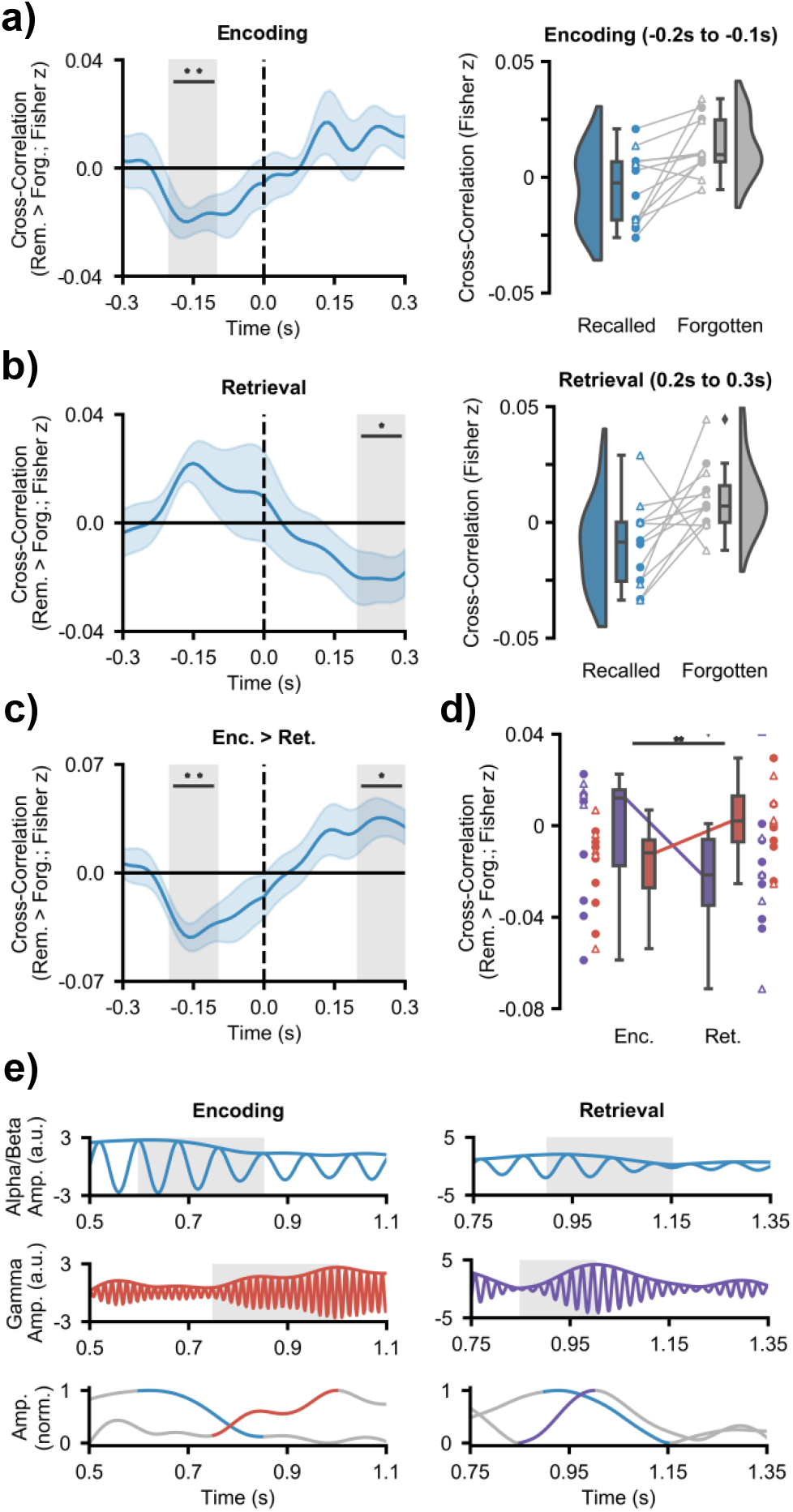
Hippocampal-neocortical time-series cross-correlations. **(a)** mean cross correlation (with shaded standard error of the mean; left) between the hippocampal “fast” gamma power and ATL alpha/beta power during encoding [**p_fdr_<0.01]. ATL power decreases precede hippocampal “fast” gamma power increases. Raincloud plot (right) depicts the difference in cross-correlation between remembered and forgotten items. Coloured circles represent participants who took part in experiment 1. Uncoloured triangles represent participants who took part in experiment 2. **(b)** mean cross correlation (with shaded standard error of the mean; left) between the hippocampal “slow” gamma power and ATL alpha/beta power during retrieval [*p_fdr_= 0.05]. Hippocampal “slow” gamma power increases precede ATL alpha/beta power decreases. Raincloud plot (right) depicts the difference in cross-correlation between remembered and forgotten items. Coloured circles represent participants who took part in experiment 1. Uncoloured triangles represent participants who took part in experiment 2. **(c)** the contrast of cross-correlation activity between encoding and retrieval [*p_fdr_ < 0.05,**p_fdr_<0.01]. **(d)** Mean cross-correlation between neocortical alpha/beta power and hippocampal gamma power (“slow” in purple; “fast” in red; with standard error of the mean) as a function of memory operation (top: subject level; bottom: electrode-pair level). A repeated-measures ANOVA reveals an interaction between hippocampal gamma frequency and memory task when predicting memory-related hippocampal-neocortical cross-correlation (**p < 0.01). **(e)** filtered single trial traces at encoding (left) and retrieval (right), in the ATL (top) and hippocampus (middle). The envelopes of these traces are plotted beneath. During encoding, a reduction in ATL alpha/beta activity precedes an increase in hippocampal “fast” gamma power. During retrieval, an increase in hippocampal “slow” gamma power precedes a decrease in ATL alpha/beta activity.

### Hippocampal gamma power increases and neocortical alpha/beta power decreases cooperate during the encoding and retrieval of human episodic memories

So far, we have demonstrated that both neocortical alpha/beta power decreases and hippocampal fast and slow gamma power increases arise during episodic memory processes. Critically however, the synchronisation/desynchronisation framework^3^ would predict that these two markers correlate in such way that neocortical power decreases precede hippocampal power increases during encoding while hippocampal power increases precede neocortical power decreases during retrieval. Such a hypothesis can be tested through the use of cross-correlation, where the time-series of neocortical alpha/beta power is offset relative to the time-series of hippocampal gamma power in an attempt to identify at what time lag the two time-series most strongly correlate. A negative lag indicates that early neocortical signals correlate with late hippocampal signals, while a positive lag indicates that early hippocampal signals correlate with late neocortical signals. Like traditional correlations, a negative correlation (from here termed ‘anticorrelation’) indicates an increase in one metric is accompanied by a decrease in the other.

At encoding, we hypothesised that the degree of neocortical power decreases can predict the degree of hippocampal gamma power increases (i.e. a negative lag anticorrelation). On a cognitive level, this would signify information processing within the neocortex preceding representational binding in the hippocampus. The cross-correlation was computed for every trial, and the memory-related difference was calculated by subtracting the mean cross-correlation across forgotten items from the mean cross-correlation across remembered trials. By calculating the memory-related difference, any correlation between the two time-series that is driven by shared noise (originating from a shared reference) is removed, as this reference-related correlation is consistent across remembered and forgotten trials (additional analysis in the SI appendix confirms that shared reference activity does not account the observed effects reported here). Furthermore, the memory-related difference highlights memory-specific dynamics in neocortical-hippocampal links, rather than general, memory-unspecific connectivity. In line with our hypothesis, later remembered items showed a significant anticorrelation at a negative lag between ATL alpha/beta power and hippocampal “fast” gamma power relative to later forgotten items (p_fdr_ = 0.006, d = 0.961; see fig. 4a for difference line plot). This cross-correlation suggests that alpha/beta power decreases precede “fast” gamma power increases by approximately 100-200ms. No correlation was observed between ATL alpha/beta power and hippocampal “slow” gamma power at any lag. These results indicate that a unique connection exists between the ATL and the hippocampus during episodic memory formation, where ATL power decreases precedes hippocampal “fast” gamma power increases.

We then investigated whether this relationship reverses during episodic memory retrieval (i.e. hippocampal power increases precedes neocortical power decreases). On a cognitive level, this would represent pattern completion in the hippocampus preceding information reinstatement in the neocortex. To test this, we repeated the cross-correlation analysis in the same manner as above for epochs covering the presentation of the retrieval cue and then calculated the memory-related difference by subtracting the mean cross-correlation across forgotten items from the mean cross-correlation across remembered trials. Relative to forgotten items, remembered items showed a significant anticorrelation at a positive lag between ATL alpha/beta power and hippocampal “slow” gamma power (p_fdr_ = 0.037, d = 0.731; see fig. 4b), where an increase in hippocampal gamma power preceded a decrease in ATL alpha/beta power by 200-300ms). No correlation was observed between ATL alpha/beta power and hippocampal “fast” gamma power at any lag. These results indicate that hippocampal “slow” gamma power increases precede ATL alpha/beta power decreases during the retrieval of episodic memories – a reversal of the dynamic observed during episodic memory formation.

We then examined how the neocortical-hippocampal dynamics differed between encoding and retrieval. To this end, the subsequent memory effect (SME; remembered minus forgotten cross-correlation at encoding) for ATL alpha/beta power and hippocampal “fast” gamma power was contrasted with the retrieval success effect (RSE; remembered minus forgotten cross-correlation at retrieval) for ATL alpha/beta power and hippocampal “slow” gamma power in a group level, non-parametric, permutation test. This revealed an interaction whereby ATL power decreases preceded hippocampal power increases during encoding (p_fdr_ = 0.005, d = 1.181; 100-200ms) but hippocampal power increases preceded ATL power decreases during retrieval (p_fdr_ = 0.025, d = 0.855; 200-300ms) [see fig. 4c]. These results support those reported in the previous two paragraphs; 1) ATL alpha/beta power decreases precede hippocampal “fast” gamma power increases during episodic memory formation and 2) hippocampal “slow” gamma power increases precedes ATL alpha/beta power decreases during episodic memory retrieval.

Lastly, we examined whether the “fast” gamma effect was specific to encoding and the “slow” gamma effect was specific to retrieval. To this end, we conducted a non-parametric, permutation-based, 2×2 repeated measures ANOVA (memory operation × gamma frequency), taking encoding-related activity from the −200 to −100ms time bin and retrieval-related activity from the 200 to 300ms time bin. Analysis revealed a significant interaction between the two factors (p = 0.001; partial η^2^ = 0.172). The interaction (as pictured in Figure 4d) suggests that the hippocampal “fast” gamma power negatively cross-correlated with ATL alpha/beta power to a greater degree than hippocampal “slow” gamma power during encoding, while the hippocampal “slow” gamma power negatively cross-correlated with ATL alpha/beta power to a greater degree than hippocampal “fast” gamma power during retrieval.

Notably, these effects cannot be explained by any epileptic activity such as IEDs (inter-epileptical discharges) travelling between the cortex and hippocampus. IEDs are broadband, so, one may expect that IEDs that are temporally-correlated across regions may give rise to spurious coupling between frequency bands. While certainly true, this cannot explain the effects observed here for two reasons. (1) Our findings are bidirectional – there would need to be pathological activity generated in both the ATL and the hippocampus to produce such bidirectional hippocampal-cortical interactions, where IEDs generated in the ATL travel to the hippocampus to produce the encoding effect, and IEDs generated in the hippocampus travel to the ATL produce the retrieval effect. None of the patients who took part in the experiment had pathological tissue in both the ATL and the hippocampus, so the IED confound explanation cannot explain the directionality of our effect. (2) IEDs are broadband, yet our effects are narrowband. During encoding, we observe the cross-correlation between neocortical alpha/beta and hippocampal fast gamma, but importantly not neocortical alpha/beta and hippocampal slow gamma. Any IED-induced broadband artifact would inherently yield cross-correlations with alpha/beta power and both gamma bands, and not within one singular band. Complementary qualitative analysis to support this conclusion can be found in the supplementary information.

## Discussion

To successfully encode and recall episodic memories, we must be capable of 1) representing detailed multisensory information, and 2) binding this information into a coherent episode. Numerous studies have suggested that these two processes are accomplished by neocortical desynchronisation (as measured by decreases in oscillatory power) and hippocampal synchronisation (as measured by increases in fast and slow oscillatory gamma power) respectively^3,5,7,26^. Here, we provide the first empirical evidence that these two processes co-exist and interact. During successful episodic memory formation, alpha/beta power decreases in the anterior temporal lobe (ATL) reliably precede “fast” hippocampal gamma power increases (60-80Hz) by 100-200ms. In contrast, “slow” hippocampal gamma power increases (40-50Hz) precede alpha/beta power decreases by 200-300ms during successful episodic memory retrieval. These findings demonstrate that the interaction between neocortical alpha/beta power decreases and hippocampal power increases in distinct, functionally-relevant gamma rhythms underpins the formation and retrieval of episodic memories.

Our central finding demonstrates that ATL alpha/beta power decreases and hippocampal fast and slow gamma power increases interact during the formation and retrieval of episodic memories, respectively. This result draws together a multitude of conflicting studies, some of which indicate that synchronisation benefits memory^e.g.43–45^ and others which indicate that desynchronisation benefits memory^e.g.13,24,46^, and provides a possible empirical resolution to the so-called “synchronisation-desynchronisation conundrum”^3^. These findings are in line with previous observations demonstrating that hippocampal gamma power increases precede hippocampal alpha power decreases during associative memory retrieval^47^. However, we are the first to show that this sequence reverses during encoding, and to link these two mechanisms across brain regions (via simultaneous hippocampal-neocortical recordings unavailable to ^47^). We speculate that the delay in hippocampal response relative to ATL alpha/beta power decreases during encoding reflects the need for the ATL to process semantic details prior to the hippocampus binding this information into a coherent representation of the event^26,27^. In contrast, we posit that the ATL delay in response relative to hippocampal gamma power increases during retrieval reflects the need for the hippocampal representational code to be reactivated prior to reinstating highly-detailed stimulus-specific information about the event^48^. Anatomically speaking, this reciprocal communication may be facilitated by the “direct intrahippocampal pathway” – a route with reciprocal connections between the ATL and hippocampus via the entorhinal cortex^49,50^. These anatomical connections would allow the ATL and hippocampus to cooperate during episodic memory formation and retrieval, facilitating the flow of neocortical information into the hippocampus during encoding and the propagation of hippocampal retrieval signals into the neocortex during retrieval.

We also uncovered the first empirical evidence of distinct gamma rhythms supporting human episodic memory formation and retrieval^7,35^. Specifically, we found greater “fast” gamma oscillatory activity (60-80Hz) during encoding and greater “slow” gamma oscillatory activity (40-50Hz) during retrieval, generalising earlier rodent findings ^e.g.31^ to humans. We uncovered similar distinctions in “fast” and “slow” gamma band activity when investigating memory-related changes in power and neocortical-hippocampal cross-correlations, providing additional evidence for such a distinction. Earlier rodent studies have suggested that the distinction between the two gamma bands reflects a difference in CA1 coupling^31^; “fast” gamma oscillations support CA1-entorhinal cortex coupling, facilitating the transfer of information into the hippocampus, while “slow” gamma oscillations support CA1-CA3 coupling, facilitating the reactivation of stored information. We speculate that these patterns of connectivity extrapolate to humans and explain the observed differences in gamma frequency relating to episodic memory formation and retrieval. In sum, our results suggest that “fast” and “slow” gamma activity relates to distinct processes in the successful formation and retrieval of episodic memory.

In combination, the cross-correlation and gamma-band analyses produce a detailed picture of information flow during episodic memory formation and retrieval. Based on earlier frameworks^3,7^ and models^4^, we postulate that the link between neocortical alpha/beta power decreases and hippocampal “fast” gamma power increases during memory formation reflects the flow of semantic information (processed in the ATL) to entorhinal cortex^27^ via the direct intrahippocampal pathway^49,50^, where “fast” gamma synchronicity between the entorhinal cortex and CA1 passes this information onto the hippocampus^31,51^. In contrast, the link between hippocampal “slow” gamma power increases and neocortical alpha/beta power decreases during memory retrieval reflects the flow of reactivated representational codes from CA3 to CA1 (via “slow” gamma synchronicity^31,51^), which propagates out into the neocortex^48^ via reciprocal connections in the direct intrahippocampal pathway, reinstating semantic details in the desynchronised ATL. However, future research with direct recordings from these hippocampal sub-regions in humans is needed to empirically test this proposed flow of information during episodic memory formation and retrieval.

Two questions remain however: First, do similar bi-directional streams of information flow exist between the hippocampus and other neocortical regions? As it was not medically necessary, electrode coverage did not expand to every neocortical region linked to episodic memory. Therefore, we could not test this theory. We speculate, however, that similar bi-directional links do exist. For example, hippocampal gamma power increases may interact with alpha/beta power decreases in the visual cortex to facilitate the encoding and retrieval of visual memories^20^. Speculating further, hippocampal gamma power increases may be the metaphorical spark that lights the fuse of memory replay, coded in desynchronised neocortical alpha phase patterns^19^.

Second, does the observed “fast”/”slow” gamma distinction reflect two true narrowband oscillations? While we have uncovered a distinction between “fast” and “slow” gamma frequencies during encoding and retrieval, we cannot say with certainty whether these differences are driven by two distinct oscillators, as proposed by others^31,36,52^. Indeed, one could argue that the observed differences are driven by fluctuations in the frequency of a single oscillator. While we are unaware of such a phenomenon in hippocampal gamma, such an effect has been reported in neocortical alpha^53^. Notably however, the reported alpha-band fluctuations were very subtle (<0.5Hz), so it’d be highly questionable to interpret the much larger 25Hz shift between “fast” and “slow” hippocampal power as originating from this alpha-band ‘fluctuation’ mechanism. One could alternatively argue that the width of a single oscillator frequency may fluctuate as a function of memory operation, giving an apparent shift in the ratio between “fast” and “slow” gamma. However, such an effect should introduce a symmetrical change around the peak. This is not present in our data, which suggests that such an effect is ill-suited to explain the observed difference in “fast” and “slow” gamma. In short, while any electrophysiological effect can be interpreted in many ways, it seems the most parsimonious explanation here is that distinct “fast” and “slow” gamma bands differentially influence memory operations, as proposed by Colgin^7^.

In summary, we deliver the first empirical evidence that neocortical power decreases and hippocampal power increases cooperate during the formation and retrieval of episodic memories, providing evidence that may help resolve the so-called “synchronisation-desynchronisation conundrum”^3^. Furthermore, we provide the first evidence that distinct hippocampal gamma oscillations service human episodic memory formation and retrieval, with faster (∼60-80Hz) oscillations supporting encoding and slower (∼40-50Hz) oscillations supporting retrieval. In conjunction, these results further illuminate our understanding of how interactions between the neocortex and hippocampus help build and retrieve memories of our past experiences.

## Methods

### Participants

Twelve patients (n = 8 from Queen Elizabeth Hospital Birmingham, UK; n = 4 from University Hospital Erlangen, Germany; 41.7% female; mean age = 35.5 years, range = 24 to 53 years) undergoing treatment for medication-resistant epilepsy took part in the experiment. These participants had intracranial depth electrodes implanted for diagnostic purposes. Ethical approval was granted by the NHS Health Research Authority (15/WM/0219) and the Ethik-Kommission der Friedrich-Alexander Universität Erlangen-Nürnberg (142_12 B). Informed consent was obtained in accordance with the Declaration of Helsinki.

### Behavioural paradigm: word-dynamic associative task

Seven of the twelve participants completed this paired associates task (see fig. 1c). During encoding, participants were presented with a 3 second video or sounds, followed by a word in the participant’s native language (English, n = 7; German; n = 1; presented for 3 seconds). There was a total of four videos and four sounds, repeated throughout each block. All four videos had a focus on scenery that had a temporal dynamic, while the four sounds were melodies performed on 4 distinct musical instruments. Due to time restraints, some participants only completed the experiment using one modality of dynamic stimulus (sound, n=1; video, n=5; both, n=2). Participants were asked to “vividly associate” these two stimuli. For each pairing, participants were asked to rate how plausible (1 for very implausible and 4 for very plausible) the association they created was between the two stimuli (the plausibility judgement was used to keep participants on task rather than to yield a meaningful metric). The following trial began immediately after participants provided a judgement. If a judgement was not recorded within 4 seconds, the next trial began. This stopped participants from elaborating further on imagined association they had just created. After encoding, participants completed a 2-minute distractor task which involved making odd/even judgements for random integers ranging from 1 to 99. Feedback was given after every trial. During retrieval, participants were presented with every word that was presented in the earlier encoding stage and, 3 seconds later, asked to identify the associated video/sound from a list of all four videos/sounds show during the previous encoding block. The order in which the four videos/sounds were presented was randomised across trials to avoid any stimulus-specific preparatory motor signals contaminating the epoch. Following selection, participants were asked to rate how confident they felt about their choice (1 for guess and 4 for certain). Each block consisted solely of video-word pairs or solely of sound-word pairs – there were no multimodal blocks. Each block initially consisted of 8 pairs, with each dynamic stimulus being present in two trials. However, the number of pairs increased by steps of 8 if the number of correctly recalled pairs was greater than 60% - this ensured a relatively even number of hits and misses for later analysis. Participants completed as many blocks/trials as they wished. Any participant that had fewer than 10 “remembered” or 10 “forgotten” trials after iEEG pre-processing were excluded from further analysis.

All participants completed the task on a laptop brought to their bedside. Responses were logged using the ‘f’, ‘g’, ‘h’ and ‘j’ keys, which corresponded to values ‘1’, ‘2’, ‘3’, and ‘4’. To aid comprehension, snippets of paper were placed on top of each relevant keyboard keys with the associated numerical value written upon them. The auditory stimuli were presented via the laptop’s speakers due to concerns that earphones could prove painful to the participants following electrode implantation just above the ear.

### Behavioural paradigm: animal-face-place associative task

Five of the twelve participants completed this paired associates task (see fig. 1d). During encoding, participants were first presented with an image cue of an animal for 2 seconds, followed by a pair of 2 images made up of any combination of a famous face or a famous place (i.e. face-place, face-face or place-place pairs; presented for 2 seconds). There were initially a total of 20 image pairs, repeated throughout each block. This number was reduced if the hit-rate fell below 66.25%, or increased if the hit-rate surpassed 73.75%. Participants were asked to “vividly associate” these two stimuli. For each pairing, participants were asked whether the association was plausible or implausible (the plausibility judgement was used to keep participants on task rather than to yield a meaningful metric). Participants were self-paced in providing a judgement, and the following trial began immediately afterwards. After encoding, participants completed a distractor task which involved making odd/even judgements for 15 sequentially presented random integers, ranging from 1 to 99. Feedback was given after every trial. During retrieval, participants were presented with every animal image cue that were presented in the earlier encoding stage and, 2 seconds later, asked how many of the associated face or place pairs they remember (participants had the option of responding with 0, 1 or 2). If the participant remembered at least one image, they were then asked to select the pair of images from a panel of four images shown during the previous encoding block (2 targets & 2 distractors). Participants were self-paced during the retrieval stage, though the experiment ended after a runtime of 40 minutes in total. All participants completed the task on a laptop brought to their bedside. Any participant that had fewer than 10 “remembered” or 10 “forgotten” trials after iEEG pre-processing were excluded from further analysis.

### Behavioural coding

For the first associative task, trials were classified as “remembered” if the participant selected the correct dynamic stimulus and stated that they were highly confident about their choice (i.e. scored 4 on the 4-point confidence scale). Trials were classified as “forgotten” if the participant selected the incorrect dynamic stimulus, did not respond, or stated that they guessed their choice (i.e. scored 1 on the 4-point confidence scale). For the second associative task, trials were classified as “remembered” only if the participant indicated that they remembered both images and subsequently selected both correctly from the panel. Trials were classified as “forgotten” in all other cases, where the participant indicated that they did not remember at least one image and/or subsequently selected one of the images incorrectly from the panel.

### Statistical analysis

While the two tasks differed in external stimulation, the underlying cognitive and neural phenomena relating to hypotheses is expected to be consistent across tasks. Therefore, the data for the two tasks were pooled. Unless explicitly stated otherwise in the results section, all statistics were conducted on the group level (i.e. random effects) using non-parametric, permutation based statistical tests. In analyses where multiple comparisons were made (e.g. time-series differences), the false-discovery rate correction^54^ was applied (denoted as p_fdr_). Effect sizes accompany each reported p-value; Cohen’s d was used for all t-tests (denoted as d). For reference, Cohen^55^ suggested that d=0.8 indicates a large effect, d=0.5 indicates a medium effect, and d=0.2 indicates a small effect. Partial eta squared was used as a measure of effect size for all ANOVAs (denoted as partial η^2^). For reference, partial η^2^ = 0.25 indicates a large effect, partial η^2^ = 0.09 indicates a medium effect, and partial η^2^ = 0.01 indicates a small effect.

### iEEG acquisition and preprocessing

First, the raw data was epoched; for encoding trials, epochs began 2 seconds before the onset of the visual/auditory stimulus and ended 4 seconds after verbal stimulus onset (9 seconds in total); for retrieval trials, epochs began 2 seconds before, and ended 4 seconds after, the onset of the verbal cue (6 seconds in total). Second, the data was filtered using a 0.2Hz finite-impulse response high-pass filter and 3 finite-impulse response band-pass filters at 50±1Hz, 100±1Hz and 150±1Hz, attenuating slow-drifts and line noise respectively. Third, as the iEEG data was sampled at the physician’s discretion (512Hz, n=1; 1024Hz, n=8), all data was down-sampled to 500Hz. Fourth, the data from each electrode was re-referenced to an electrode on the same shaft that was positioned in white matter (determined by visual inspection of the participant anatomy; see below). The use of a common reference electrode for both the hippocampus and neocortex ensured that any difference in electrophysiological signal from the two regions could not be explained by a difference in reference. Finally, the data was visually inspected and any trials exhibiting artefactual activity were excluded from further analysis. Any electrodes exhibiting persistent ictal and interictal activity (as identified through visual inspection) were discarded from analysis.

### Electrode localisation

First, hippocampal and white matter contacts were defined based on anatomical location through visual inspection of the T1-weighted anatomical scan (N.B. one participant had no hippocampal contacts, and therefore was excluded from all hippocampal-based analyses). Then, the native space co-ordinates of all remaining contacts were determined by visual inspection of each participant’s post-implantation T1 scan. These contact co-ordinates were then transformed from native space to MNI space using a transform matrix obtained by normalising participant T1 scans in SPM 12. These contacts were then marked as within the anterior temporal lobe (ATL) or elsewhere (this latter group was excluded from further analysis).The ATL was defined as all parts of the temporal lobe (as defined by the *wfupickatlas* plugin^56^ for SPM 12) anterior to a plane perpendicular to the long axis of the temporal lobe^57^. The plane was slightly shifted from that described in ^57^ to [y=−5, z=−30; y=15, z=−5] for the pragmatic reason of ensuring that all participants had electrode contacts in the ATL ROI. For visualisation in figure 1d, every electrode from every participant was given a diameter of 1cm and then placed in a template brain registered in MNI space. The number of electrodes in each voxel was then summed to provide a measure of summed density.

### 1/f correction

Spectral power was computed using 199 linearly-spaced 5-cycle wavelets ranging from 1 to 100Hz. The time-frequency decomposition method was kept consistent across all frequency bands to ensure that only a single slope (characterising the full extent of the 1/f dynamic) needed to be calculated and subsequently subtracted from the signal (in line with previous experiments that have extracted the 1/f characteristic from the signal ^e.g.39,40^). A vector containing values of each wavelet frequency (*A*) and another vector containing the power spectrum for each electrode-sample pair (*B*) were then log-transformed. The linear equation *Ax = B* was solved using least squares regression, where *x* is an unknown constant describing the curvature of the 1/f characteristic. The 1/f fit (*Ax*) was then subtracted from the log-transformed power-spectrum (*B*).

### Peak frequency analysis

Raw signal recorded at every contact for each epoch was convolved with a 5-cycle wavelet (0 to 1500ms post-stimulus [padded with real data for lower frequencies], in steps of 25ms; 1Hz to 100Hz, in steps of 0.5Hz). The 1/f noise was subtracted using the method described above to help pronounce the peaks in the power-spectrum. The data was then smoothed using a Gaussian kernel (full-width half-maximum 200ms; 1Hz) to attenuate inter- and intra-individual differences in spectral responses^53^ and to help approximate normally distributed data (an assumption frequently violated in small samples). The data was averaged across all time-points, trials and contacts (separately for the hippocampus and ATL). Peaks of 1/f corrected absolute power were then identified using the *findpeaks()* peak-detection algorithm implemented in Matlab (see SI appendix for details). To identify the memory-related difference in the dominant gamma bands, the power spectra for “remembered” trials were calculated in an identical manner, except that the Gaussian kernel was expanded to account for the greater variability of high-frequency oscillatory responses (200ms, 5Hz). The power-spectra for encoding and retrieval were then collapsed in seven 10Hz bins ranging from 30Hz to 100Hz and contrasted in a group level (i.e. random effects), non-parametric permutation test^58^ with 5000 randomisations. The multiple comparison issue was solved using the false-discovery rate correction ^54^. This analysis was repeated for the “forgotten” trials.

#### Selection of peak frequencies

The peak frequencies of each patient were determined using the MATLAB function *findpeaks()* on the averaged power spectrum around the approximate frequency bands (theta: 1-7Hz; alpha/beta: 8-20Hz; “slow” gamma: 30-60Hz; “fast” gamma: 50-100Hz). The bandwidths of these peaks were kept consistent across participants, and were determined through inspection of the group-averaged bandwidth of the peaks (theta: ±0.5Hz; alpha/beta: −1Hz/+5Hz [capturing the observed asymmetry in the peak]; “slow”/”fast” gamma: ±10Hz). Individual peak frequencies are reported in Supplementary Table 1.

### Spectral power analysis

For all spectral power analyses (i.e. encoding and retrieval epochs), the data underwent the same wavelet convolution, 1/f correction, and smoothing approaches described in the *peak frequency analysis* section. The data was then z-transformed using the means and standard deviations of each electrode-frequency pair^14^. The time-frequency resolved data was then averaged over electrodes of each ROI. For time-series statistical analysis, trials were split into two groups based on whether the stimuli were remembered or forgotten. Then, the time-series were collapsed into seven time bins of 200ms and the two conditions were contrasted using the same non-parametric statistical procedure described in the *peak frequency analysis* section. For statistical analyses of the interaction between memory task (encoding vs. retrieval) and gamma frequency (“fast” vs. “slow”), this memory-related difference in power (i.e. SME and RSE) was averaged over time and contrasted in a non-parametric, permutation based 2×2 repeated measures ANOVA.

### Cross-correlation analysis

For all cross-correlation analyses (i.e. encoding and retrieval epochs), the data underwent the same wavelet convolution, 1/f correction, and smoothing approaches described in the *spectral power analysis* section, with two exceptions: 1) wavelet convolution occurred in steps of 10ms rather than 50ms (enhancing temporal resolution), and 2) the temporal aspect of the smoothing kernel was reduced to 50ms to avoid excessive smoothing obscuring the temporal dynamics of the neocortical-hippocampal cross-correlation. For each “trial × electrode combination” pair, the cross-correlation between the hippocampus and the ATL, and the cross-correlation between the hippocampus and PTPR, was computed using the Matlab function crosscorr() with a lag of 300ms (meaning the correlation between hippocampus and neocortex was considered for every offset from where the neocortex preceded the hippocampus by 300ms to where the neocortex lagged behind the hippocampus by 300ms). This returned a time-series of Pearson correlation values describing the relationship between hippocampus and neocortex at all considered lags. These correlation values were then averaged over electrodes and split into two groups: remembered and forgotten. These two groups were individually averaged over trials for each participant, collapsed into bins of 100ms, and then contrasted using the same non-parametric statistical procedure described in the *peak frequency analysis* section. We term the “remembered > forgotten” difference in cross-correlation for encoding data “the subsequent memory cross-correlation” and the difference for retrieval data “the retrieval success cross-correlation”.

To test the “encoding-retrieval” × “lag-lead” difference, we contrasted the subsequent memory cross-correlation with the retrieval success using the same non-parametric statistical procedure described in the *peak frequency analysis* section.

Lastly, to test the influence of the “memory task” × “gamma frequency” interaction on the memory-related cross-correlation differences, we conducted a non-parametric, permutation-based 2×2 repeated measures ANOVA in the same manner as described in the *spectral power analysis* section.

