## Supplementary Materials for "Directional coupling of slow and fast hippocampal gamma with neocortical alpha/beta oscillations in human episodic memory"

SUPPLEMENTARY INFORMATION

### Supplementary Results

*Event-related potentials*

Measures of low-frequency oscillatory power (<10Hz) can be distorted by event-related potentials (ERPs). To rule out the concern that the observed memory-related power effects are driven by ERPs, the ERPs themselves were examined. The data was epoched from 500ms pre-stimulus to 1000ms post-stimulus and high frequency activity was filtered out using a Butterworth low-pass filter of 20Hz. Baseline correction was computed as the difference in amplitude from a pre-stimulus window (-250 to 0ms). Trials were split based on whether the stimulus pairs were remembered or forgotten, and the data was then averaged across contacts of the same region of interest. Remembered trials were contrasted with forgotten items in a non-parametric, group-level t-test. At encoding, no memory-related difference in ERPs was observed in the ATL ($p_{fdr}$ = 0.178) or hippocampus ($p_{fdr}$ = 0.368). Similarly during retrieval, no memory-related difference in ERPs was observed in the ATL ($p_{fdr}$ = 0.291) or hippocampus ($p_{fdr}$ = 0.279).

*The influence of reference on cross-correlation results*

The choice of reference can have a drastic impact on cross-regional analysis such as the cross-correlation result reported in the main text. Indeed, it is possible that as neocortical and hippocampal electrodes on the same shaft shared a common (white matter) reference, an artificial correlation between two regions may arise as a result of shared reference activity. However, such a correlation for power can only be positive and zero centred which is in stark contrast to the negative non-zero centred correlation reported in the main text. In the main text, this potential confound was tackled by contrasting hits with misses to subtract out any reference-related confound (as both conditions should fall foul to the issue) while preserving statistical power. Here, we complimented the subtraction approach by reproducing the cross-correlation analysis using different references for hippocampal and neocortical electrodes. The analytical approach is identical to that reported in the main text with one key exception: when creating hippocampal-neocortical electrode pairs, any pair from the same shaft (i.e. shared the same white matter reference) was excluded from analysis (leaving only electrode pairs with different white matter reference). Notably, one participant had only a single shaft where the hippocampus and ATL was recorded from, meaning that all hippocampal and ATL recordings in this participant shared a white matter reference. This participant was therefore excluded from this analysis. This approach yielded near identical results to those reported in the main text. ATL alpha/beta power decreases preceded hippocampal fast gamma power increases during successful memory formation ($p_{fdr}$ = 0.026, d = 0.954), while hippocampal slow gamma power increases preceded ATL alpha/beta power decreases during successful memory retrieval ($p_{fdr}$ = 0.035, d = 0.741). Similarly, the 2 x 2 (gamma frequency x encoding/retrieval) was replicated (p = 0.013, partial eta squared = 0.136). These findings suggest that the cross-correlation results are not due to spurious correlations owing to the choice of referencing.

*The influence of interictal discharges on cross-correlation results*

Notably, epileptic activity such as IEDs (Inter-Epileptical Discharges) can travel between the neocortex and the hippocampus, and these discharges can influence whether a memory is encoded/retrieved or not. While the data was cleaned of these discharges, a control analysis was run where any electrodes near the seizure foci (as described in supplementary table 4) were excluded. It is important to note

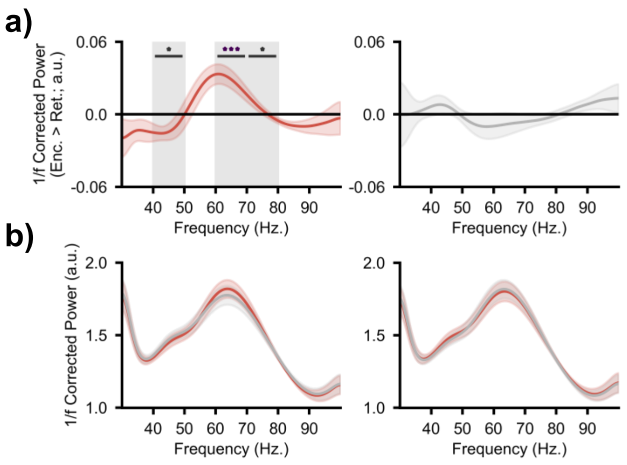


**Supplementary Figure 1.** Encoding-retrieval differences in hippocampal gamma. **(a)** the difference in encoding hippocampal gamma power and retrieval hippocampal gamma power for remembered items (left) and forgotten items (right). **(b)** the power spectra for encoding and retrieval hippocampal gamma power for remembered items (left) and forgotten items (right). A clear "fast" gamma peak can be seen between 60 and 70Hz. A second notch can be seen between 40 and 50Hz, reflecting "slow" gamma activity.

that only seven subjects maintained electrodes in both the ATL and the hippocampus, and therefore were included in this control analysis.

For this approach, the memory-related alpha-gamma contrast in the ATL at encoding and at retrieval continued to demonstrate a statistically significant effect ($p_{fdr}$ = 0.041, Cohen's d = 0.485, and $p_{fdr}$ = 0.036, Cohen's d = 1.302). Similarly, the encoding-retrieval contrast within the ATL matched those reported in the main text (-100 to -200ms, $p_{fdr} < 0.001$, Cohen's d = 1.104; 200 to 300ms, $p_{fdr} < 0.001$, Cohen's d = 1.647). Lastly, the 2 x 2 (gamma frequency x encoding/retrieval) continued to reveal a significant interaction (p = 0.036, partial eta squared = 0.234). Together, these results analytically demonstrate that interictal discharges cannot explain the observed results.

### Supplementary Discussion

*On the "fast"/"slow" gamma distinction:* Notably, the raw power spectra (see supplementary figure 1b) do not show two clear gamma peaks, but rather a singular peak around the "fast" gamma frequency, with a smaller "slow" gamma notch on the rising slope of the gamma peak. While this may speak against the idea that two distinct gamma band oscillators differentially influence encoding and retrieval, such an effect has been reported in rodents before. Bragin and colleagues[1] demonstrated that "fast" gamma oscillations can effectively mask "slow" gamma oscillations, and the "fast" oscillator must be lesioned to observe the "slow" gamma oscillations. While lesioning would be highly unethical in humans, directly contrasting the power spectra at encoding and retrieval produces a synonymous result because it subtracts the amplitude imbalance in "fast" and "slow" gamma power common across all trials and returns the residual difference (see figure 2b). In other words, the gamma-band raw power spectra can be misleading due to masking of "slow" gamma by high-amplitude "fast" gamma, but 'encoding versus retrieval' contrasts address these issues.

**Supplementary Table 1.** Resonating frequencies of each patient (1-7 took part in task 1; 8-12 took part in task 2).

| | Neocort. Alpha/Beta | Hipp. "Slow" Gamma | Hipp. "Fast" Gamma |
|---|---|---|---|
| Patient 1 | 12.5 | 39.5 | 64.5 |
| Patient 2 | 10.0 | 38.0 | 56.0 |
| Patient 3 | 11.5 | 49.0 | 63.0 |
| Patient 4 | 12.0 | 44.5 | 66.0 |
| Patient 5 | 14.5 | 44.5 | 67.0 |
| Patient 6 | 16.0 | 46.5 | 68.0 |
| Patient 7 | 10.0 | 45.5 | 70.0 |
| Patient 8 | 10.5 | 47.5 | 71.0 |
| Patient 9 | 10.0 | 40.0 | 56.0 |
| Patient 10 | 10.5 | 44.5 | 63.5 |
| Patient 11 | 10.5 | 47.0 | 60.5 |
| Patient 12 | 10.0 | 38.5 | 64.5 |
| **Mean** | **11.5** | **43.8** | **64.2** |

**Supplementary Table 2.** Number of electrodes in each region of interest per participant (1-7 took part in task 1; 8-12 took part in task 2).

| | Hippocampus | Anterior Temporal Lobe | Total |
|---|---|---|---|
| Patient 1 | 3 | 3 | 6 |
| Patient 2 | 3 | 3 | 6 |
| Patient 3 | 1 | 4 | 5 |
| Patient 4 | 4 | 3 | 7 |
| Patient 5 | 1 | 3 | 4 |
| Patient 6 | 2 | 3 | 5 |
| Patient 7 | 2 | 3 | 5 |
| Patient 8 | 3 | 2 | 5 |
| Patient 9 | 2 | 2 | 4 |
| Patient 10 | 2 | 3 | 5 |
| Patient 11 | 2 | 7 | 9 |
| Patient 12 | 2 | 3 | 5 |
| **Total** | **27** | **39** | **66** |
| **Median** | **2** | **3** | **5** |

**Supplementary Table 3.** Number of trials per condition, per participant following artefact rejection (1-7 took part in task 1; 8-12 took part in task 2).

| | Encoding | | | Retrieval | | |
|---|---|---|---|---|---|---|
| | Remembered | Forgotten | Total | Remembered | Forgotten | Total |
| Patient 1 | 14 | 48 | 62 | 15 | 62 | 77 |
| Patient 2 | 19 | 19 | 38 | 22 | 26 | 48 |
| Patient 3 | 30 | 18 | 48 | 35 | 21 | 56 |
| Patient 4 | 31 | 26 | 57 | 37 | 24 | 61 |
| Patient 5 | 31 | 29 | 60 | 36 | 31 | 67 |
| Patient 6 | 90 | 52 | 142 | 91 | 51 | 142 |
| Patient 7 | 21 | 12 | 33 | 21 | 16 | 37 |
| **Mean** | **33.7** | **29.1** | **62.9** | **36.7** | **33.0** | **69.7** |
| Patient 8 | 63 | 94 | 157 | 69 | 93 | 162 |
| Patient 9 | 28 | 22 | 50 | 45 | 28 | 73 |
| Patient 10 | 113 | 32 | 165 | 144 | 40 | 184 |
| Patient 11 | 173 | 48 | 221 | 186 | 18 | 204 |
| Patient 12 | 147 | 70 | 217 | 145 | 69 | 214 |
| **Mean** | **104.8** | **53.2** | **162.0** | **117.8** | **49.6** | **167.4** |

**Supplementary Table 4.** Additional patient demographics (1-7 took part in task 1; 8-12 took part in task 2).

| Patient No. | Years of Seizures | Dominant Hand | Epileptic Foci |
|---|---|---|---|
| Patient 1 | 17 | Right | Left hippocampus |
| Patient 2 | 3-5 | Right | Bilateral medial temporal lobe |
| Patient 3 | 34 | Unknown | Left frontal lobe |
| Patient 4 | 9-12 | Right | Bilateral medial temporal lobe |
| Patient 5 | 8 | Right | Right neocortical temporal lobe |
| Patient 6 | 10-12 | Left | Right medial parietotemporal lobe |
| Patient 7 | 26 | Right | Right medial temporal lobe |
| Patient 8 | Unknown | Right | Unknown |
| Patient 9 | 26 | Right | Right medial temporal lobe |
| Patient 10 | Unknown | Right | Unknown |
| Patient 11 | Unknown | Left | Left frontal lobe |
| Patient 12 | 22 | Right | Left medial temporal lobe |
